# Evolution of a biological thermocouple by adaptation of cytochrome c oxidase in a subterrestrial metazoan

**DOI:** 10.1101/2023.12.05.570156

**Authors:** Megan N. Guerin, TreVaughn Ellis, Mark J. Ware, Alexandra Manning, Ariana Coley, Ali Amini, George Chung, Kristin C. Gunsalus, John R. Bracht

## Abstract

In this study we report a naturally evolved temperature-sensing electrical regulator in the cytochrome c oxidase of the Devil Worm, *Halicephalobus mephisto.* This extremophile metazoan was isolated 1.3 km underground in a South African goldmine, where it adapted to heat and potentially to hypoxia, making its mitochondrial sequence a likely target of adaptational change. We obtained the full mitochondrial genome sequence of this organism, and show through dN/dS analysis statistically robust evidence of positive selection in *H. mephisto* cytochrome c oxidase subunits. Seventeen of these positively-selected amino acid substitutions were localized in proximity to the H- and K-pathway proton channels of the complex. Surprisingly, the *H. mephisto* cytochrome c oxidase proton pump completely shuts down at low temperatures (20°C) leading to approximately a 4.8-fold reduction in the transmembrane proton gradient voltage (ΔΨ_m_) compared to optimal temperature (37°C). Direct measurement of oxygen consumption found a corresponding 4.7-fold drop at 20°C compared to 37°C. Correspondingly, the lifecycle of *H. mephisto* takes four-fold longer at the low temperature compared to higher. This elegant evolutionary adaptation creates a finely-tuned mitochondrial temperature sensor, allowing this ectothermic organism to maximize its reproductive success in varying environmental temperatures. Our study shows that evolutionary innovation may remodel core metabolism to make it more accurately map onto environmental variation.

## Introduction

*Halicephalobus mephisto*, or the Devil Worm, is a subterranean nematode isolated from 1.3 kilometers below Earth’s surface ^1^. Endemic to mine water that is warm (37℃), hypoxic (0.42–2.3 mg/L dissolved O_2_,), alkaline (pH 7.9), and methane-rich ^1^, the nuclear genome displayed expanded novel stress response gene families including heat-shock protein (Hsp70) and avrRpt2 induced gene 1 (AIG1)^2^. Here we report the full mitochondrial genome sequence of *H. mephisto* and report the first analysis of cytochrome c oxidase function in this unique organism.

The mitochondria is a vital, double-membraned organelle that is most well characterized for its role in ATP production, β-oxidation of fatty acids, and the initiation of apoptosis, while its dysfunction has been linked to human diseases including neurodegenerative disorders^3^, cancer^4^, and even autoimmune disease^5^. The human and *Drosophila* mitochondrial genomes encode 13 protein-coding genes, 22 transfer RNAs (tRNAs), and two ribosomal RNAs (rRNAs), while most nematodes have reduced their protein-coding gene repertoire to 12^6^.

The evolution of mitochondrial genomes remains poorly understood, though given their asexual reproduction, lack of recombination, and highly polyploid nature, the selection pressures acting on mitochondria are quite different from those involving nuclear genes^7,8^. Mitochondrially encoded genes, and COX1 specifically, are under particularly strong functional constraint for performing electron transport and ATP generation and thus, are under potent purifying selection^7,9–11^. Here we show that evolution can extensively remodel the COX1 sequence to make mitochondrial function exquisitely temperature-selective. Because this modification of cytochrome c oxidase creates a temperature-sensing electrical voltage (proton gradient), it functions as a naturally-evolved thermocouple device.

## Results

PacBio long-read sequencing of the *H. mephisto* mitochondrial genome produced a 14,349 bp sequence that was 81% AT-rich, including 12 protein-coding genes, 22 tRNAs, and two rRNAs, that agreed closely with those of other nematodes including those of the *Halicephalobus* genus (Figure 1). Along with most nematodes, *H. mephisto* has lost the *atp8* gene, and all genes are transcribed in the same direction, a characteristic of Chromadorea^6^. The control region is 863 bp long, located between tRNA-Isoleucine and tRNA-Arginine, and a remarkable 95% AT (Figure 1). Of mitochondrial genomes sequenced to date, *H. mephisto* encodes one of the most AT-rich.

**Figure 1.**
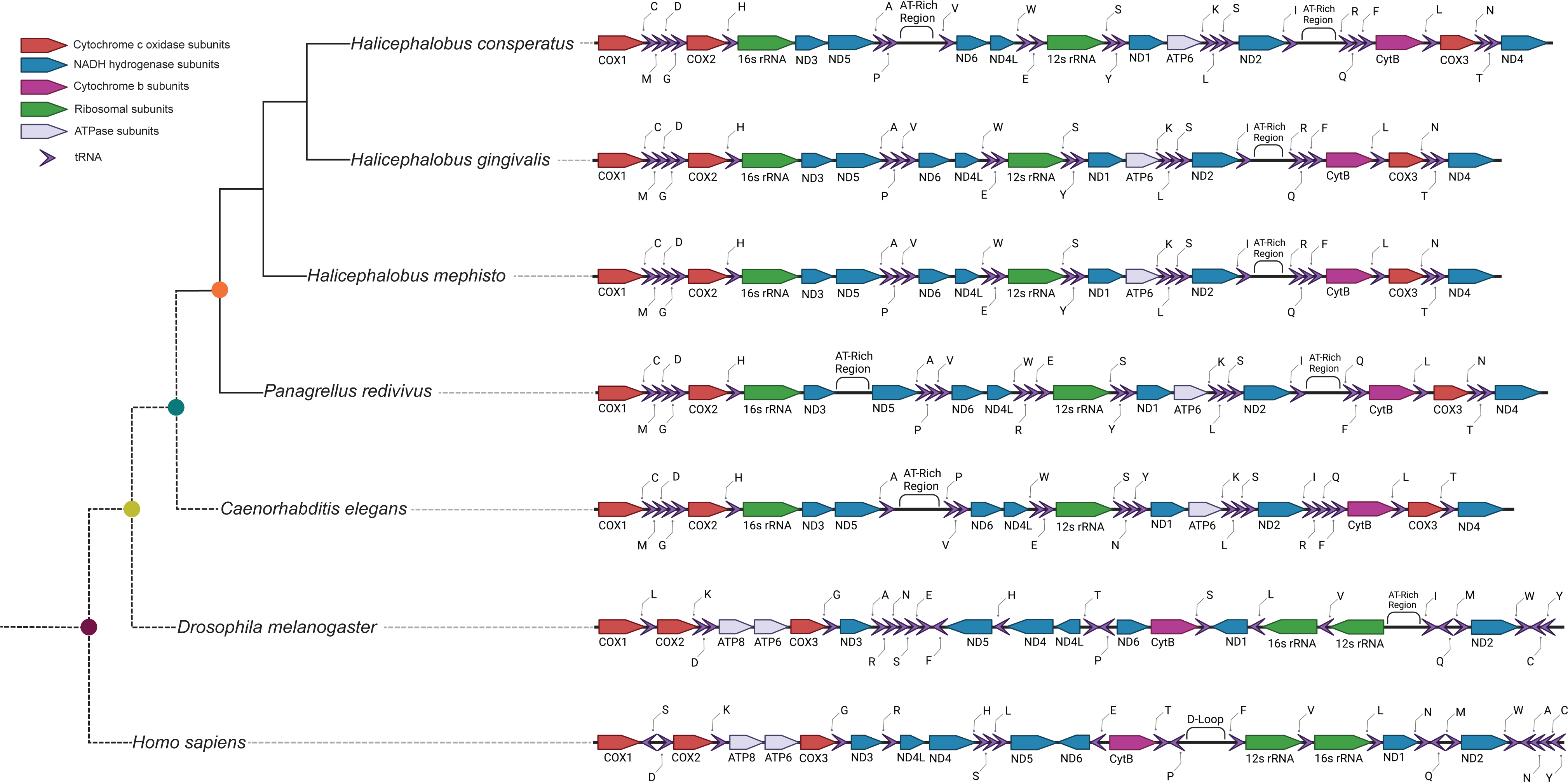
Overview of mitochondrial architectures in human, *Drosophila,* and nematodes. Note, *Halicephalobus consperatus* was formerly *Halicephalobus* strain NKZ332. Transfer RNAs indicataed by their corresponding single-letter amino acid.

To broaden our analysis we included mitochondrial sequences of two additional *Halicephalobus* species: *H. gingivalis,* a horse parasite^12^, and species NKZ332, originally isolated by Natsumi Kanzaki in association with the Japanese termite, *Reticulitermes speratus* and sequenced by Erik Ragsdale^13^. Both the nuclear and mitochondrial genomes of this organism show distinct molecular markers from all known *Halicephalobus* species, and in phylogenetic analysis this species is a well resolved sister group to both *H. mephisto* and *H. gingivalis.* Therefore we have renamed strain NKZ332 *Halicephalobus consperatus* (for ‘with *speratus*’) after its association with the termite. Unfortunately the nature of the interaction, whether phoretic, symbiotic, or parasitic, may never be resolved because the species is no longer in culture (E. Ragsdale, personal communication). For phylogenetic analysis we included 35 additional mitochondrial genomes sampling broadly across nematode phylogeny, and include two outgroups: *Homo sapiens,* and *Drosophila melanogaster* (accession numbers in Supplemental Table 1).

Utilizing the 12 protein-coding genes of these 38 mitochondrial genomes we constructed a concatenated mitochondrial protein ML tree (Figure 2a) which is broadly consistent with other nematode mitochondrial gene trees^14^. In particular, the monophyly of classes Enoplia (Clade II) and Chromadorea (containing Clades III, IV, and V) are cleanly recovered; however within Chromadorea both clades III (Spirurina) and IV (Tylenchina) are not monophyletic, consistent with previous mitochondrial phylogenies^6,14^. Clade V (Rhabditina) is monophyletic with the exception of two newly placed *Diploscapter* species (*D. pachys* and *D. coronatus*) emerging as sister clades to *Bursaphelenchus xylophilus* (Clade IV) (Figure 2a). Given that Maximum Likelihood analysis may suffer from long-branch attraction artifacts^15^ we re-analyzed our data by Bayesian methods, recovering a nearly identical tree again with both *Diploscapter* species emerging as sisters to *B. xylophilus* (Figure S1). Given the abundance of nuclear genomic data demonstrating that both *Diploscapter* species are closer relatives of *C. elegans* and members of Rhabditina^16,17^, not Tylenchina, we conclude that the mitochondrial genomes of these species exhibit significant homoplasy, but we have not explored this phenomenon further.

**Figure 2.**
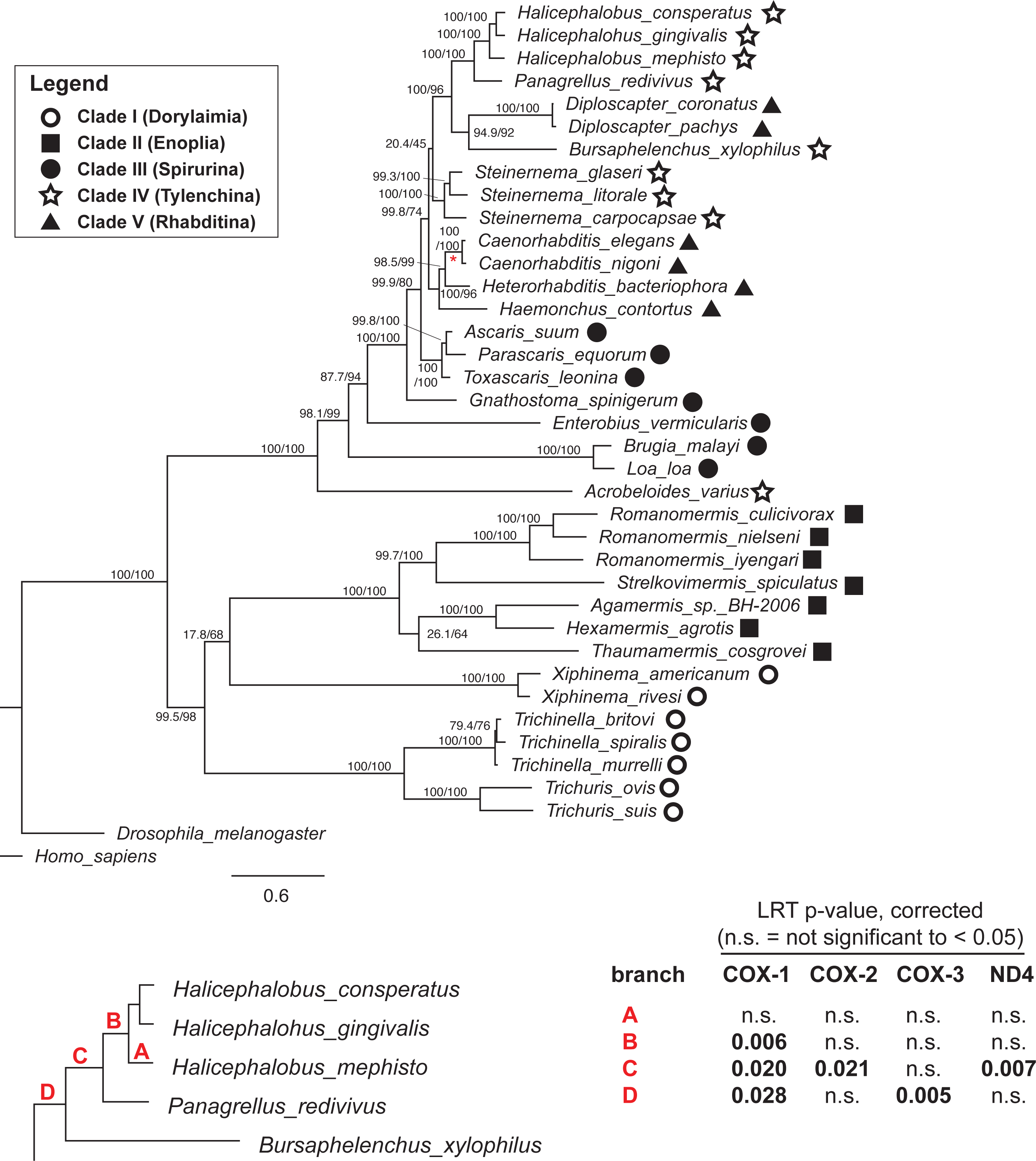
Phylogeny and dN/dS analysis. **a**, Combined mitochondrial ML phylogeny of nematodes created with IQtree and model mtZOA+F+I+G4. Bootstrap values in parentheses indicate SH-aLRT support (%) / ultrafast bootstrap support (%). Scale bar indicates substitutions per site. **b**, branch labels and Paml analysis by branch-sites. In the table the p-values for the Likehood Ratio Tests (LRTs) for positive selection are shown. For each branch PAML was run twice, once to allow an estimated dN/dS (ω2) for a subset of sites on the foreground branch; once to fix ω2 at 1 for the foreground branch (in all cases another ω1, measures purifying selection of most sites). By taking the ratio of maximum likelihood values, a p-value is measured using a Chi-Square table with 2 degrees of freedom (critical values are 5.991 for p-value < 0.05 and 9.210, p-value < 0.01). Bonferroni correction was performed to correct for multiple hypothesis testing.

To evaluate potential selective events in the *Halicephalobus* lineage we performed a likelihood ratio test (LRT) of positive selection using PAML^18^, which requires the phylogeny to accurately capture the species’ evolutionary history. Therefore, we manually re-located the two *Diploscapter* species to their correct position (based on nuclear phylogenetics) within Rhabditina as indicated by a red asterisk on Figure 2a. Supporting the correctness of this move, the PAML-derived log-likelihood (lnL) values increased by an average of 23.2 across the branches shown in Figure 2b after relocating *Diploscapter* (max, 25.2 and min, 22.3). We tested COX1, 2, and 3 and ND4 using the branch-sites test for positive selection (also known as Model A), generating Bonferroni-corrected p-values for each branch^19^. While we had expected positive selection might be detectable along the most recent lineage leading to *H. mephisto* (branch A), we instead found statistically robust positive selection along three ancestral branches B, C, and D (Figure 2b). COX1 is a particular focus of evolutionary innovation as the only gene with positive selection across three of four branches tested (Figure 2b).

The division of branches into A, B, C, and D divides the evolutionary selection into distinct times: the divergence of the suborder Tylenchina (branch D); the diversification of family Panagrolaimidae (branch C); the common *Halicephalobus* ancestor (branch B); finally branch A leading to *H. mephisto*. We therefore analyzed amino acid substitution patterns in COX1 along these lineages. To do this, we examined fixed derived non-synonymous mutations (FdNs) shared within the clades ^20^ (Table 1). We identified 18 amino acid substitutions in COX1, one in COX2, and one in COX3 by manual inspection of sequence alignments, for 20 total substitutions (Table 1). In each case a single nonsynonymous single nucleotide polymorphism (snp) was responsible for the substitution and was conserved within the clade; sometimes with adjacent synonymous sequence changes preserving the altered amino acid (Table 1). Given the wide range of evolutionary divergence between species within Tylenchina, Panagrolamidae, and even within *Halicephalobus* (see below), this pattern of derived amino acid substitutions suggests a functional preservation and is consistent with positive selection^20^.

PAML also reports sites of selection as part of its output. These sites we interpret with caution given the high divergence between sequences in our phylogeny (see discussion) raising concerns around synonymous site (dS) saturation noted in some tests of positive selection^21^. The PAML sites corroborated the alignment-based FdNs sites but also identifies other sites. For example, for COX1, PAML identified 3 sites on branch B (1 is an FdNs); 14 sites on branch C (9 FdNs); and 12 sites on branch D (8 are FdNs). Thus these PAML-identified sites are confirmatory of the FdNs identified through sequence alignment. We also note that in general the branch-sites test (which we implemented) has been found to be conservative under synonymous site saturation, displaying a loss of power rather than false positive inferences, at saturation of dS ^21^. Overall, the statistically robust inference of positive selection by LRT and 20 FdNs amino acid substitutions warrant further investigation into their function.

We hypothesized that the 20 amino acid substitutions might co-localize in the cytochrome c oxidase structure. To evaluate this, we created a structural model of the COX1, -2, and -3 proteins of *H. mephisto* by SWISS-MODEL based upon the well resolved bovine cytochrome c oxidase (3abm in PDB; 1.95Å resolution). In spite of the relatively high protein divergence (identity of *H. mephisto* to bovine proteins: COX1, 58%, COX2, 38% and COX3, 42%) a structure largely superimposable with the cow structure (Figure S2) was generated (RMSD of 0.142Å to template 3abm) (Figure 3a, b). Supporting the accuracy of the model, the amino acids lining the predicted heme binding pocket of COX1 were more conserved than the overall protein, at 81% identity, compared to 58% for the whole protein, and the bovine heme groups could be shown nestled within a pocket of the *H. mephisto* predicted structure (Figure 3a-d)

**Figure 3.**
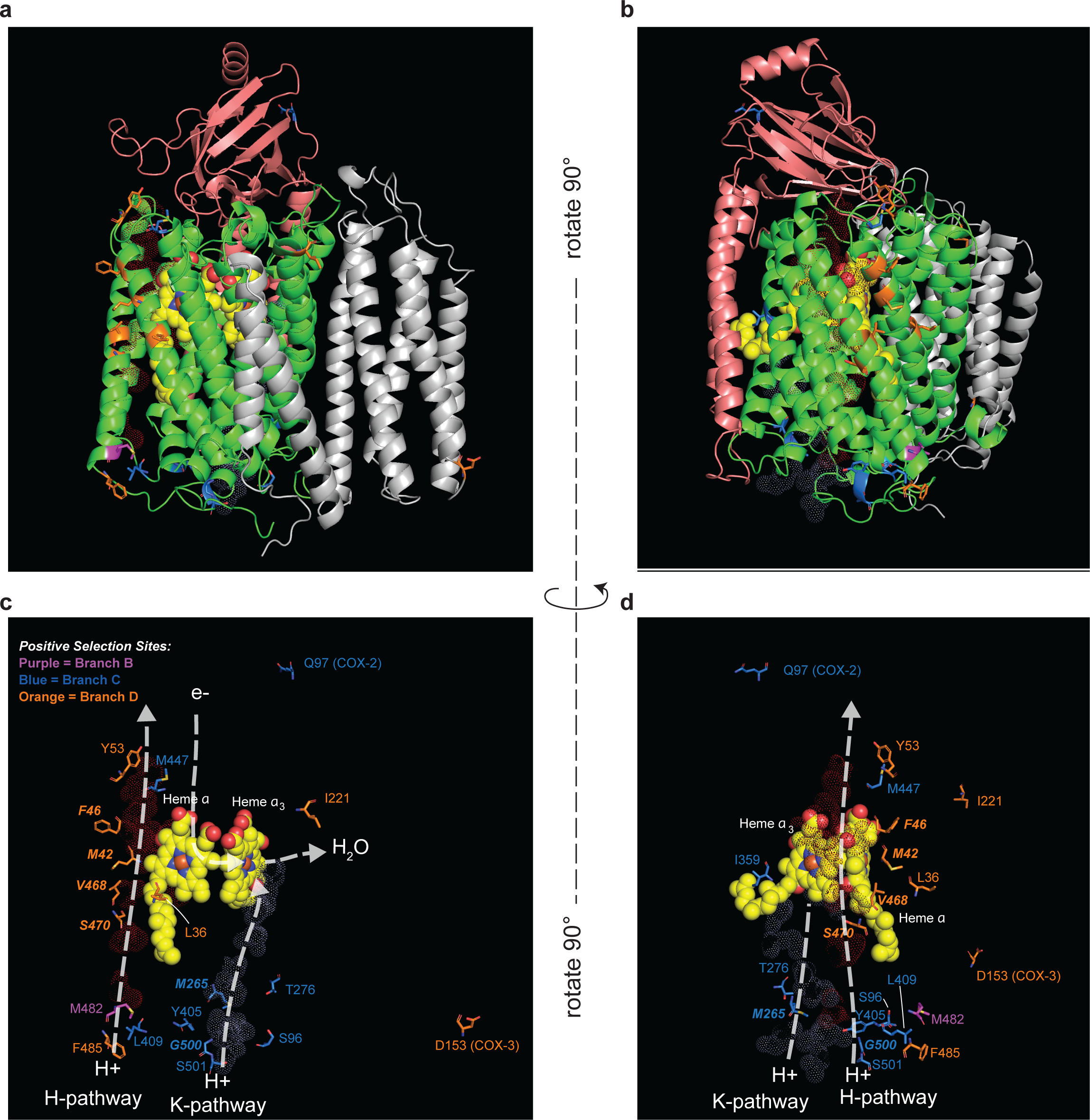
Homology model of *H. mephisto* COX1, COX2, and COX3 modeled on bovine crystal structure 3abm. **a**, ribbon structure showing the three *H. mephisto* proteins COX1 = green; COX2= salmon pink; and COX3=light gray. Heme groups from bovine crystal structure in yellow. **b**, Same as A, but rotated 90 degrees. **c**, Cutaway view showing only the heme and positively selected sites with their residue numbers. For residues within 4 Å of their channel are in bold italic font while those within 13 Å are in regular font. The H- and K-proton pathways are shown as red or light blue dots outlining the residues from the bovine structure corresponding to these pathways. **d**, Same as C but rotated 90 degrees.

We then mapped the identified FdNs amino acid changes onto this structure. Of the 18 located in COX1, 17 (94%) are proximal to (within 13 Å) proton channels: the H-pathway or the K-pathway (Figure 3c,d). Six FdNs are within 4 Å of a proton channel (Figure 3c, d, Table 1). Because of the separation of evolutionary timeframes by branch, we can see that the modification of channels occurred in stages, first along the H-pathway (branch D) then along the K-pathway (branch C) and finally, back along the H-pathway again (Branch B) (Figure 3c, d).

COX2 and -3 showed a single FdNs each, consistent with their much smaller sizes relative to COX1. For COX2 the substitution occurs on a loop near the cytochrome c binding site while for COX3 the substitution is located on the mitochondrial matrix side of the mitochondrial membrane (Figure 3a). The possible functional impact of these alterations is not clear from these data, but the COX3 substitution is very close to the interface with the COX6A2 subunit interface.

Because cytochrome c oxidase is the site of oxygen reduction to produce water, we wondered whether the observed amino acid changes might provide adaptive advantage under hypoxic conditions, particularly because *H. mephisto* was originally isolated from hypoxic subterrestrial water^1^. To test this, we performed anaerobic chamber culture using oxygen absorbing sachets at 20°C and 37°C, producing an environment of less than 0.1% oxygen. For both *H. mephisto* and *C. elegans* synchronized L1 hatching worms were cultured on standard OP50 food on NGM plates. We found that both *C. elegans* and *H. mephisto* survived 9 days of severe hypoxia at 20°C (Figure 4). However by day 22 only *C. elegans* had survivors and by 29 days all worms had perished in the hypoxic environment (Figure 4). At 37°C *H. mephisto* were markedly hypoxia intolerant, exhibiting 100% lethality by 2 days (Figure 4). Together these findings suggest that *H. mephisto* is not well adapted to severe hypoxia, and we conclude that the amino acid changes in cytochrome c oxidase are most likely involved in thermal adaptation.

**Figure 4.**
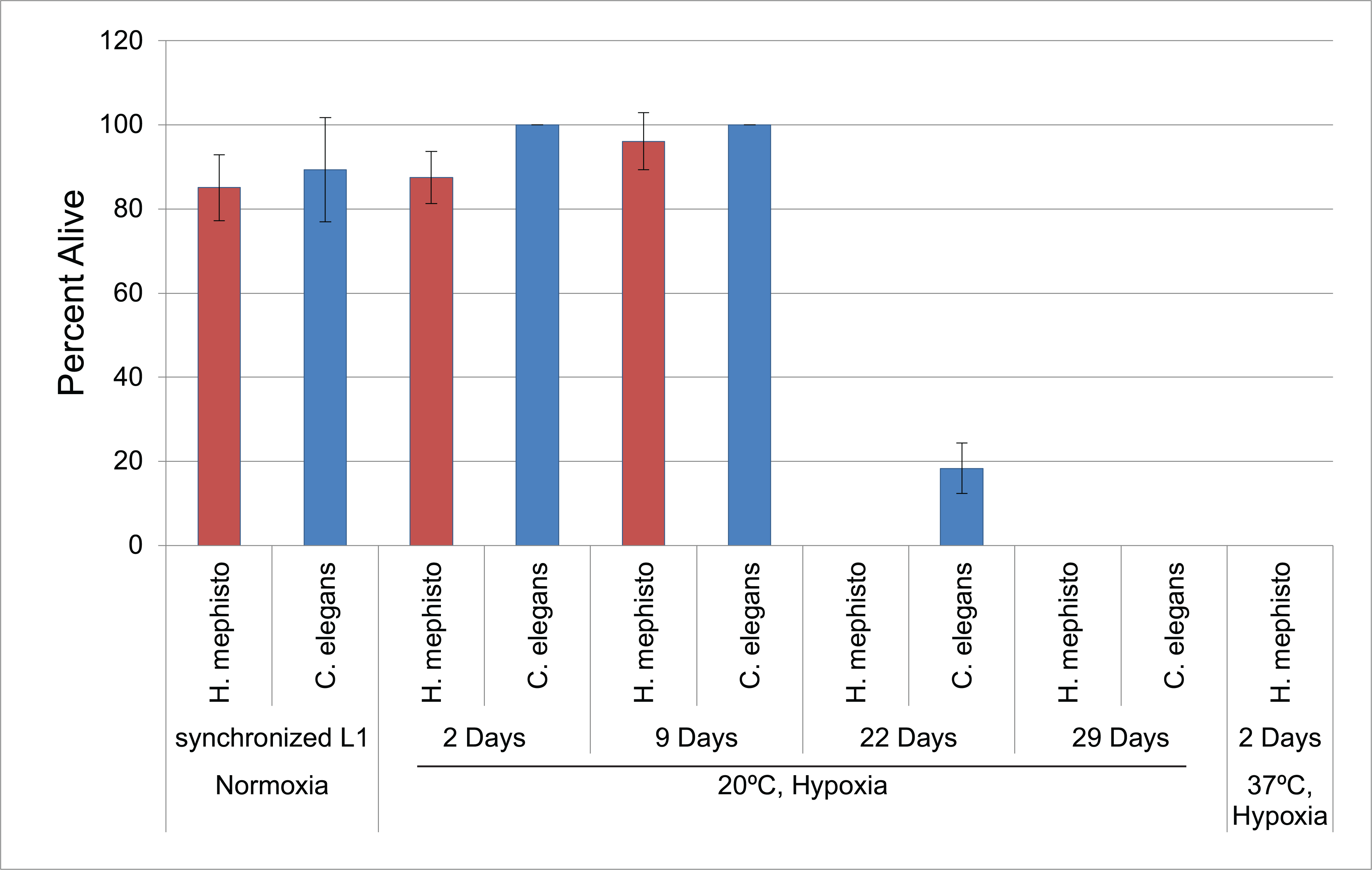
Survival of *H. mephisto* and *C. elegans* in hypoxic conditions at different temperatures. Plotted are mean and standard deviations across replicates.

Given the potentially function-altering changes to the proton translocation H- and K-pathways (Figure 3c, d) we performed analysis of the inner-membrane proton gradients using tetramethylrhodamine, ethyl ester (TMRE) (Figure 5a-e). This cationic dye is imported into the mitochondrial inner lumen proportionally to the proton gradient across the inner mitochondrial membrane, ΔΨ_m_. The proton gradient is a product of three proton pumps: complex I, III, and IV ^22^ (Figure 5f). To isolate the effect of complex IV alone, we used 25mM of sodium azide, a cytochrome c oxidase (Complex IV)-specific inhibitor, which leaves Complexes I and III unaffected^23^ (Figure 5g). Thus, by comparing untreated with azide-treated nematodes we can directly assess the contribution of Complex IV activity to the ΔΨ_m_. The terminal electron acceptor under sodium azide exposure is presumed to be fumarate rather than O_2_, as in mammalian hypoxia^24^ (Figure 5g).

**Figure 5.**
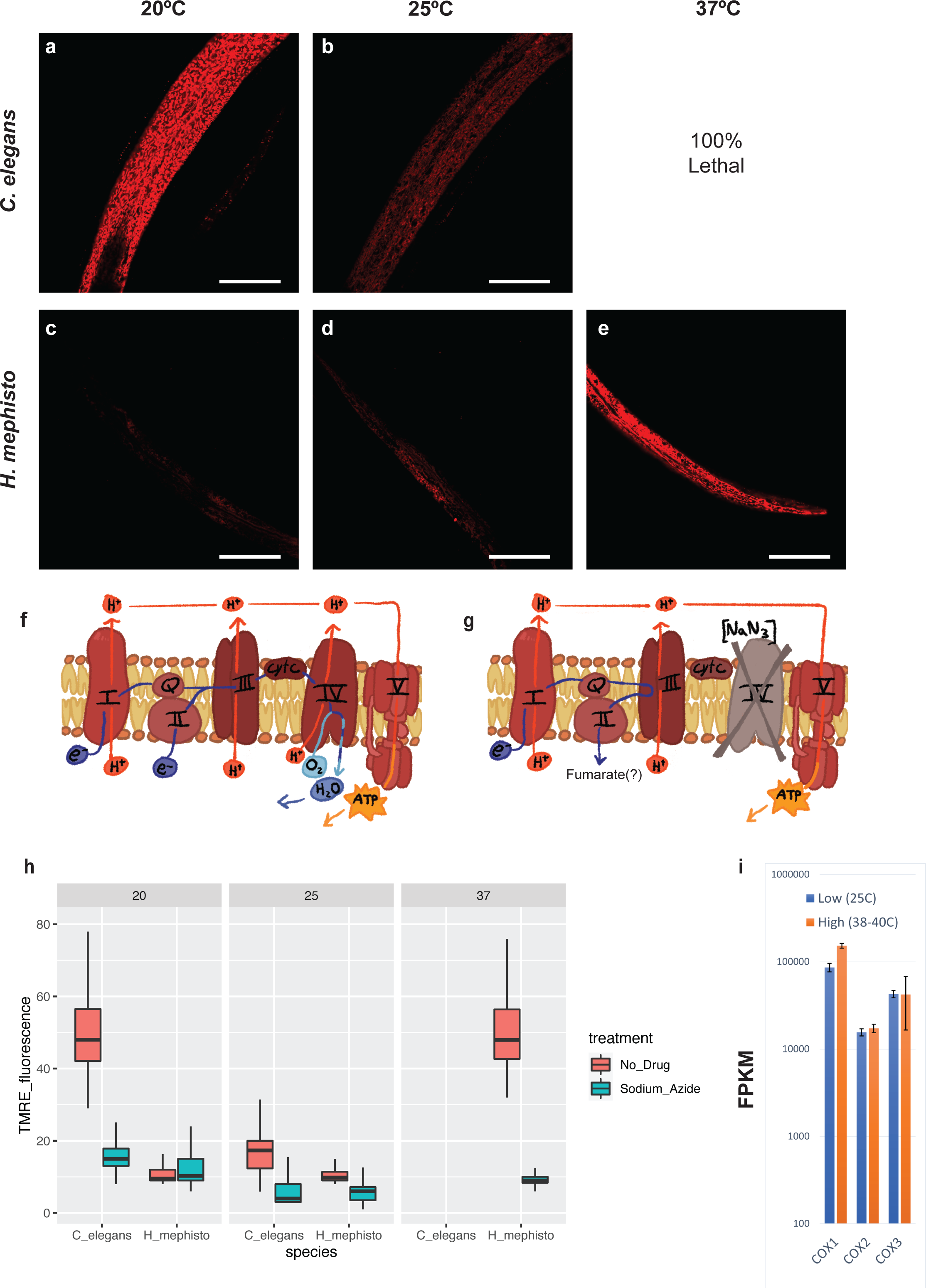
Quantification of mitochondrial proton pumping activity in *H. mephisto* and *C. elegans*. **a**, *C. elegans* at 20°C. **b**, *C. elegans* at 25°C. **c**, *H. mephisto* at 20°C. **d**, *H. mephisto* at 25°C. **e**, *H. mephisto* at 37°C. **f**, Schematic of the electron transport chain showing proton pump and electron flow. **g**, Schematic of the electron transport chain illustrating the effect of sodium azide, which blocks Complex IV while leaving remaining proton pumping Complexes I and III unaffected. **h**, Boxplot showing the relative TMRE signal from *C. elegans* or *H. mephisto* at different temperatures with or without sodium azide. All boxes in the plot are statistically different from each other, by ANOVA and Tukey’s HSD post-hoc test to p<0.001, except for the 20°C *H. mephisto* no-drug vs. sodium azide treated(p=0.99), *H. mephisto* 37°C no-drug vs. *C. elegans* 20°C no-drug (p=0.13), and 25°C *H. mephisto* sodium azide vs. *C. elegans* 25°C sodium azide (p=0.39). **i**, RNA-seq analysis of COX1, 2, and 3 expression under different temperatures. FPKM, Fragments per Kb per Million mapped reads. TMRE, TetraMethylRhodamine, Ethyl ester, perchlorate. All scale bars in worm images represent 50μm.

Using this TMRE-based imaging method, we found that both *H. mephisto* and *C. elegans* display similar ΔΨ_m_ (p=0.13, two-way ANOVA and Tukey’s HSD post-hoc test) when comparing their optimum temperatures: 20°C for *C. elegans* and 37°C for *H. mephisto* (matching its native subterrestrial habitat^1^). For *C. elegans,* at 25°C (near its thermal maximum temperature of 26°C) a significant reduction in ΔΨ_m_ is apparent, an apparent proton gradient decoupling due to thermal stress^25^. Most surprisingly, we find that at 20°C Complex IV is essentially inactive in *H. mephisto,* because sodium azide treatment makes no difference in ΔΨ_m_ (p=0.99, ANOVA and Tukey’s HSD post-hoc test, Figure 5h). These results suggest that, for *H. mephisto,* the proton pumping at 20°C is conducted largely by complexes I and III leading to a very strong reduction in the proton gradient, and thus a lower voltage, across the membrane. We observed that ΔΨ_m_ changes by 4.8-fold (Figure 5h) between 37°C and 20°C. This difference is directly due to altered cytochrome c oxidase activity: at 37°C the inhibition of cytochrome c oxidase by sodium azide led to a 5.1-fold drop in proton pumping while it only caused a 0.98-fold drop at 20°C because of the inactivity of cytochrome c oxidase at this temperature (Figure 5h).

The temperature-controlled difference in proton pumping by cytochrome c oxidase could be due to amino acid changes or to changes in gene expression. To test this, we examined temperature-dependent mitochondrial gene expression from our previously published RNAseq dataset^2^. For COX1, 2, and 3, temperature made only a slight difference in expression (Figure 5i). COX1 was elevated 1.8-fold at 38-40°C relative to 25°C while COX2 and COX3 were essentially unchanged, 1.1 and 0.98 - fold different, respectively (Figure 5i). Analysis of all 12 protein-coding mitochondrial sequences revealed that all were regulated by less than 2-fold by the temperature changes (Figure S3). Given that ΔΨ_m_ changes by 4.8-fold at 25°C relative to 37°C (ΔΨ_m_ at 20°C and 25°C are indistinguishable, Figure 5h) we conclude that the dramatic effect of high-vs-low temperature on ΔΨ_m_ cannot be explained by transcriptional regulation alone and most likely reflects a difference in the protein function, not regulation.

Mitochondrial respiration couples electron transport to its acceptor, oxygen, with the production of a proton gradient (ΔΨ_m_) and ATP. Therefore, we directly assayed the mitochondrial oxygen consumption rate with a Seahorse XF Mini analyzer^26,27^. This technique allows us to compare mitochondrial oxygen consumption in *H. mephisto* at 20°C or 37°C (Figure 6a) and *C. elegans* at 20°C (Figure 6b). The function of cytochrome c oxidase can be assessed directly by comparing untreated OCR with the sodium azide-treated OCR (OCR_untreated_ - OCR_sodium azide_). This analysis shows that cytochrome c oxidase activity is decreased by 4.7-fold at 20°C compared to 37°C, consistent with the proton gradient data (Figure 6c).

**Figure 6.**
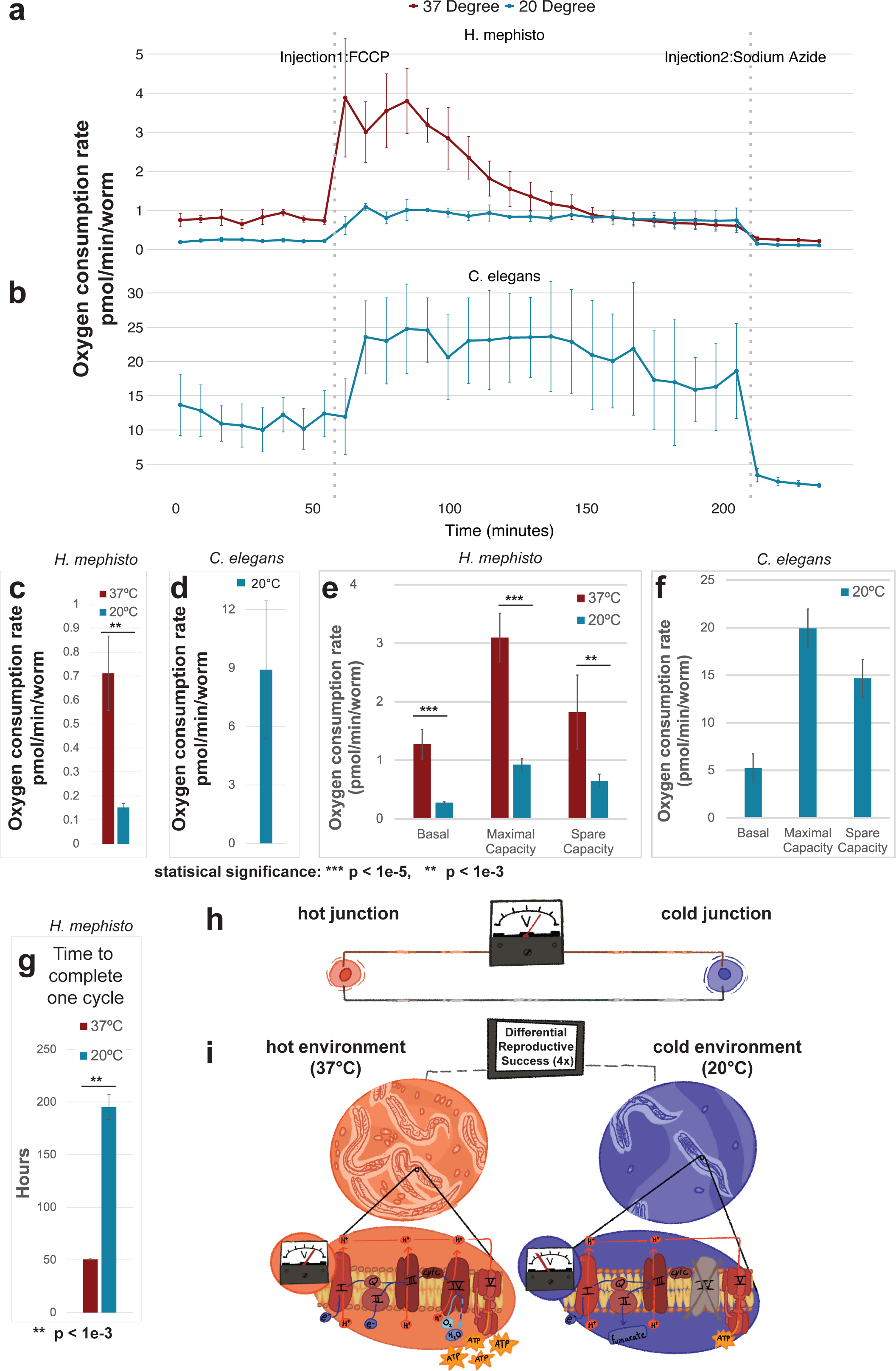
Direct measurement of oxygen consumption rates by Seahorse XF Mini. **a**, Oxygen consumption rate (OCR) for *H. mephisto* at 20°C (blue line) and 37°C (red line). Shown is mean with standard deviation of 32-point time course with 40μM FCCP and 25mM sodium azide added at the labeled dotted gray lines. **b**, Oxygen consumption rate (OCR) for *C. elegans*, plotted as in a with mean and standard deviation of 32-point time course. **c**, Cytochrome c oxidase activity (calculated as OCR_untreated_ - OCR_sodium azide_) in *H. mephisto* at 20°C and 37°C. Shown is the mean and standard error of the mean. Statistical significance assessed by Student’s two-tailed T-test for samples of unequal variance. **d**, Cytochrome c oxidase activity (calculated as OCR_untreated_ - OCR_sodium azide_) in *C. elegans* at 20°C. Shown is the mean and standard error of the mean. **e**, Basal respiration (at 1% DMSO), maximal capacity, and spare capacity for *H. mephisto* at 20°C and 37°C. Shown is the mean and standard error of the mean. Statistical significance assessed by pairwise comparisons using Wilcoxon rank sum exact test. **f**, Basal respiration (at 1% DMSO), maximal capacity, and spare capacity for *C. elegans* at 20°C. Shown is the mean and standard error of the mean. **g**, Time required (in hours) to complete one life cycle in *H. mephisto* at 20°C and 37°C. Shown are mean and standard deviation. Statistical significance assessed by Student’s two-tailed T-test for samples of unequal variance. **h**, Schematic of an engineered thermocouple device, which senses temperature as an electrical differential (voltage) between two junctions (hot and cold). **i**, Schematic of a biological thermocouple, in which varying environmental temperatures translate into both voltage differences (within the mitochondrial inner membranes) and differential reproductive success across environments.

Overall, *C. elegans* has a much higher per-worm activity of cytochrome c oxidase (Figure 6d) than *H. mephsto* (Figure 6c), which is consistent with previous *C. elegans* estimates^26,27^. By using cyanide-p-trifluoromethoxy-phenylhydrazone (FCCP), a potent decoupling agent of the inner mitochondrial membrane, the Seahorse instrument enables the calculation of basal, maximal, and spare respiratory capacity^26^. In our hands, FCCP was only efficacious in 1% DMSO (see methods) so we used 1% DMSO controls as a baseline. (In our hands 1% DMSO alone induced a statistical increase in OCR of *H. mephisto* (Figure S4a) while causing a decrease of OCR in *C. elegans* (Figure S4b).) Our data show that in *H. mephisto,* basal respiration (at 1% DMSO), maximal respiration, and spare capacity are all statistically reduced at 20°C relative to 37°C (Figure 6e) and that they are all greater in *C. elegans* (Figure 6f).

Temperature appears to play a central role in regulating *H. mephisto* metabolism.Given that at 20°C relative to 37°C the contribution of cytochrome c oxidase to OCR was reduced 4.7-fold and mitochondrial proton gradient decreases 4.8-fold we would predict these organisms experience decreased ATP production^28^, and lower metabolic rates. To test this, we measured the time per lifecycle (L1 to L1) as a proxy for metabolic rate at the two temperatures (20°C and 37°C). We found that there is a statistically significant 4-fold increase in time-per-cycle at 20°C (mean 195.2 hr) relative to 37°C (mean 50.5 hr) (Figure 6g).

## Discussion

Our analysis of COX1, COX2, and COX3 uncovered a striking evolutionary pattern of fixed, derived nonsynonymous substitutions (FdNs) along the H- and K-pathway proton translocation channels (Figure 3c, d). These substitutions are extremely likely to alter the function of cytochrome c oxidase, and several of the sites with substitutions have been identified in other systems as either critical or adjacent to critical residues in proton translocation. The H-pathway transports protons from the mitochondrial lumen through a water-accessible channel leading to heme *a,* and from there protons move through a hydrogen-bond network to be ejected into the intermembrane space^29^. A peptide bond between S441 and Y440 transfers the proton, and a S441P substitution abolishes proton pumping in HeLa cells^30^. S441 in cow corresponds to *H. mephisto* M447, one of the FdNs from Branch C, making this a particularly intriguing substitution potentially impacting proton pumping.

One FdNs, F46 (cow A39) on Branch D, is immediately adjacent to a critical residue: bovine and *H. mephisto* R38^29^. R38 acts on the opposite end of the hydrogen-bond network from cow S441/ *H. mephisto* M447, serving to feed protons from the heme *a* into the hydrogen-bond network^29,30^. (The R38 residue is completely conserved in all species we analyzed). There are 7 FdNs along the water-channel leading up to the heme *a* (F485, L409, M482, S470, L36, V468, M42) and Y53 is near the ejection site on the intermembrane space (Figure 3c, 3d). Thus the conserved nonsynonymous changes along the H-pathway go all the way from the water channel at the mitochondrial lumen to the heme-to-hydrogen-bond network and to the proton ejection site near the intermembrane space^29–31^ (Figure 3c, 3d).

Many of the substitutions make chemically impactful changes. Of the seven substitutions along the H-pathway channel, three are from nonpolar to polar and two go in the reverse (polar to nonpolar) (Table 1). On branch D, four of eight substitutions make a polar to nonpolar change or vice versa. The nine substitutions along branch C are more conservative with only three nonpolar to polar changes (Table 1).

Mitochondrially-encoded proteins are traditionally found to be under strong purifying selection and high functional constraint^7,9–11^. Supporting this, deleterious mutations are rapidly eliminated in oocytes^32^ of mice^33^, worms^34^ and flies^35^, and improved quality control of somatic deleterious mitochondrial mutations was able to extend lifespan of flies^36^. In spite of this strongly purifying pressure, notable examples of positive selection in mitochondrial genes are known, including in bats^37^, geese^38^, turtles^39^, snakes^40^, and penguins^41^.

The study of snakes found 23 amino acid sites under positive selection in COX1^40^. Of these 23 amino acid sites, three overlap with those from our study: bovine / *H. mephisto* L35 / M42, A89 / S96, and L353 / I359. Of these, only one has a the same substitution in *H. mephisto* and snakes: bovine L35 is methionine 42 in branch D nematode sequences (Table 1), and also in a blind snake species *Amerotyphlops reticulatus*; other snakes mostly have isoleucine at that position^40^. The other two sites, cow A89 / *H. mephisto* S96 and cow L353 / *H. mephisto* I359 are not convergently mutated in snakes and nematodes. The other studies, from bats^37^, geese^38^, turtles^39^, and penguins^41^, focus either on different components of the electron transport chain entirely or identify substitutions not shared in our study.

Our data suggest that *H. mephisto* adaptations are driven by adaptation to temperature, not hypoxia (Figure 4). Interestingly, combining high temperature with severe hypoxia resulted in complete lethality of *H. mephisto* after only two days (Figure 4). This synthetic lethality may not be surprising given that both hypoxia and temperature increase the production of reactive oxygen species (ROS) in human cell culture and other organisms^25,42,43^.

Our data show that *H. mephisto* metabolism is exquisitely tuned to temperature. The mitochondrial membrane proton gradient ΔΨ_m_ (Figure 5h) and oxygen consumption rate (Figure 6c) both decrease by near-identical amounts at lower temperature (4.8-fold and 4.7-fold) while lifecycle time increases by 4-fold (Figure 6g). Together these data elegantly link environmental temperature, mitochondrial respiration, and organismal reproduction.

Our data do not directly prove that positively selected residues drive the observed thermal tuning. However, several lines of evidence support a central role of the positively selected amino acid substitutions. We show that *H. mephisto* is not well-adapted to severe hypoxia (Figure 4) suggesting that positive selection is for thermal adaptation instead. Within COX1, 17 of 18 positively-selected amino acid changes co-localize to proton translocation channels, with six occurring within four angstroms of a channel. These changes would be predicted to change proton pumping and our direct experimental assessment of proton pumping confirms a striking change relative to *C. elegans* (Figure 5h). These findings were corroborated by measurements of oxygen consumption rates by cytochrome c oxidase (Figure 6c) conclusively showing that the loss of proton gradient at low temperatures is not due to increased proton leakage, which might otherwise confound our results; it is instead attributable to decreased cytochrome c oxidase activity. Combined with our data showing gene expression changes are insufficient to explain the observed temperature-driven functional differences (Figure 5i), the most plausible explanation is that the positively-selected amino acid changes contribute to a thermally-tuned mitochondrial system. However, formal proof that the 20 FdNs directly alter mitochondrial respiration awaits mitochondrial gene sequence replacement methods in either *H. mephisto* or *C. elegans* or heterologous expression studies in other organisms such as yeast. These methods largely remain to be developed.

Our work also uncovers a novel type of regulation similar to engineered thermocouples, which are electrical devices designed to detect temperature differences and convert them into voltage^44,45^. In a thermocouple device, the difference in temperature between a hot and a cold junction produces an electrical voltage (Figure 6h); in *H. mephisto* the voltage differences occur within animals distributed across the environment where they may encounter low temperatures (low voltage) or high temperatures (high voltage) (Figure 6i). An ectothermic animal would gain significant advantage from efficiently coupling temperature to reproductive rates, thereby minimizing wasteful reproduction at non-optimal thermal environments. The direct linkage of mitochondrial respiration with temperature and lifecycle represents an elegant solution to this problem, a biologically evolved thermal-couple (Figure 6i).

*C. elegans* appears to lack a thermal coupling, with no change in oxygen consumption rate observed at different temperatures^27^. Together our work demonstrates a novel type of evolved adaptational metabolic regulation occurring most likely through amino acid changes to core metabolic machinery. Adaptation may proceed through the invention of more precise mappings of metabolic and reproductive functions onto the environment.

## Acknowledgements

We acknowledge Dr. Stefano Costanzi for his guidance and suggestions regarding 3D structural modeling and visualization. We also thank our anonymous reviewers for their feedback.

## Author contributions

**Guerin, Megan** performed formal analysis, visualization, and writing-original draft

**Ellis, TreVaughn** performed formal analysis, visualization, and investigation

**Ware, Mark** performed formal analysis and visualization

**Chung, George** performed investigation (the *D. pachys* mitochondrial genome)

**Gunsalus, Kristin** performed conceptualization

**Bracht, John** performed conceptualization, visualization, formal analysis, investigation, funding acquisition, supervision, project administration, and writing-review and editing

## Data and materials availability

The mitochondrial sequence and annotation of *H. mephisto* has been deposited to GenBank under accession #OP965539.

---

**Figure S1.**
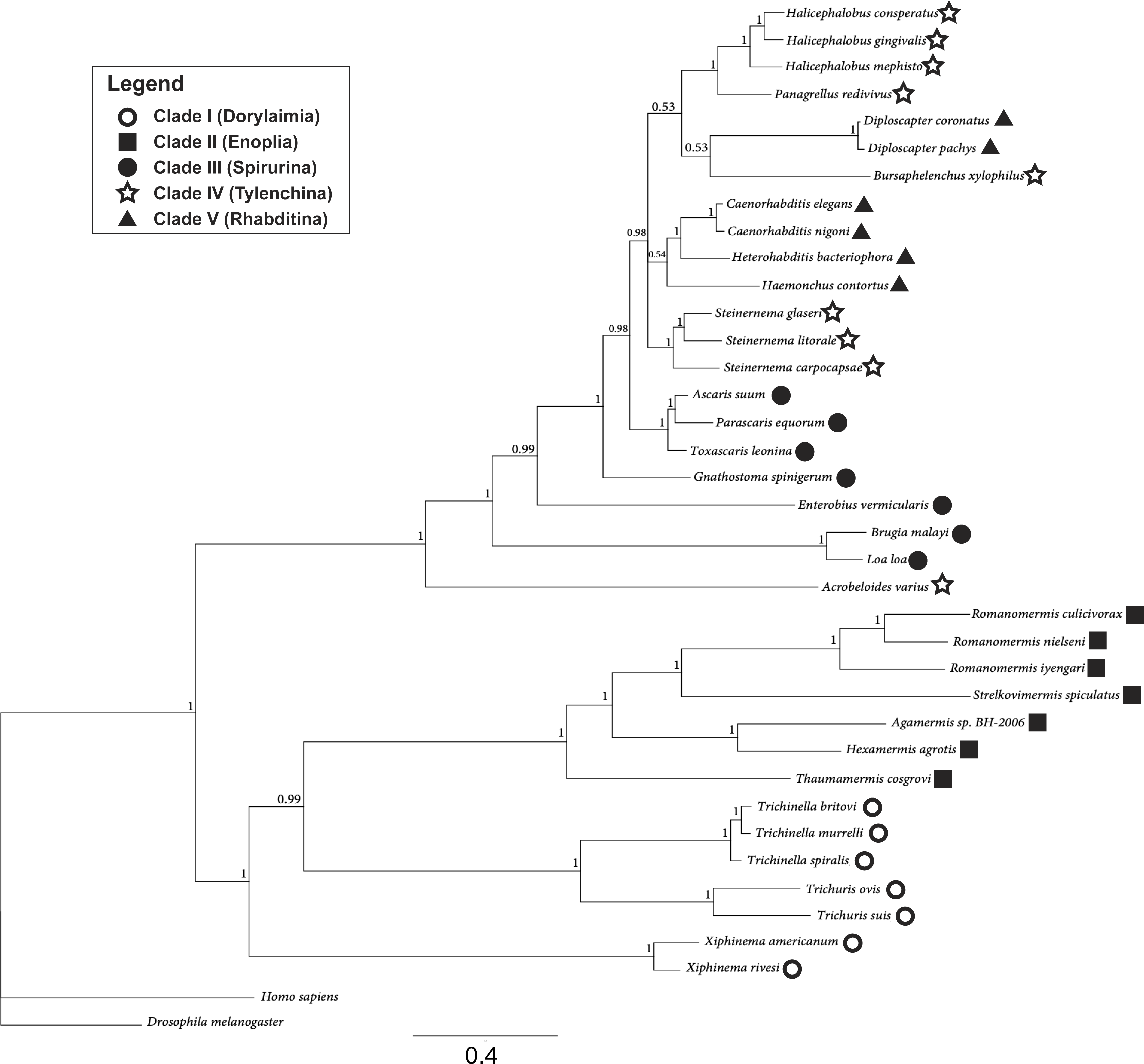
Bayesian catenated mitochondrial protein tree. Branches indicate posterior probabilities. Scale bar indicates substitutions per site.

**Figure S2.**
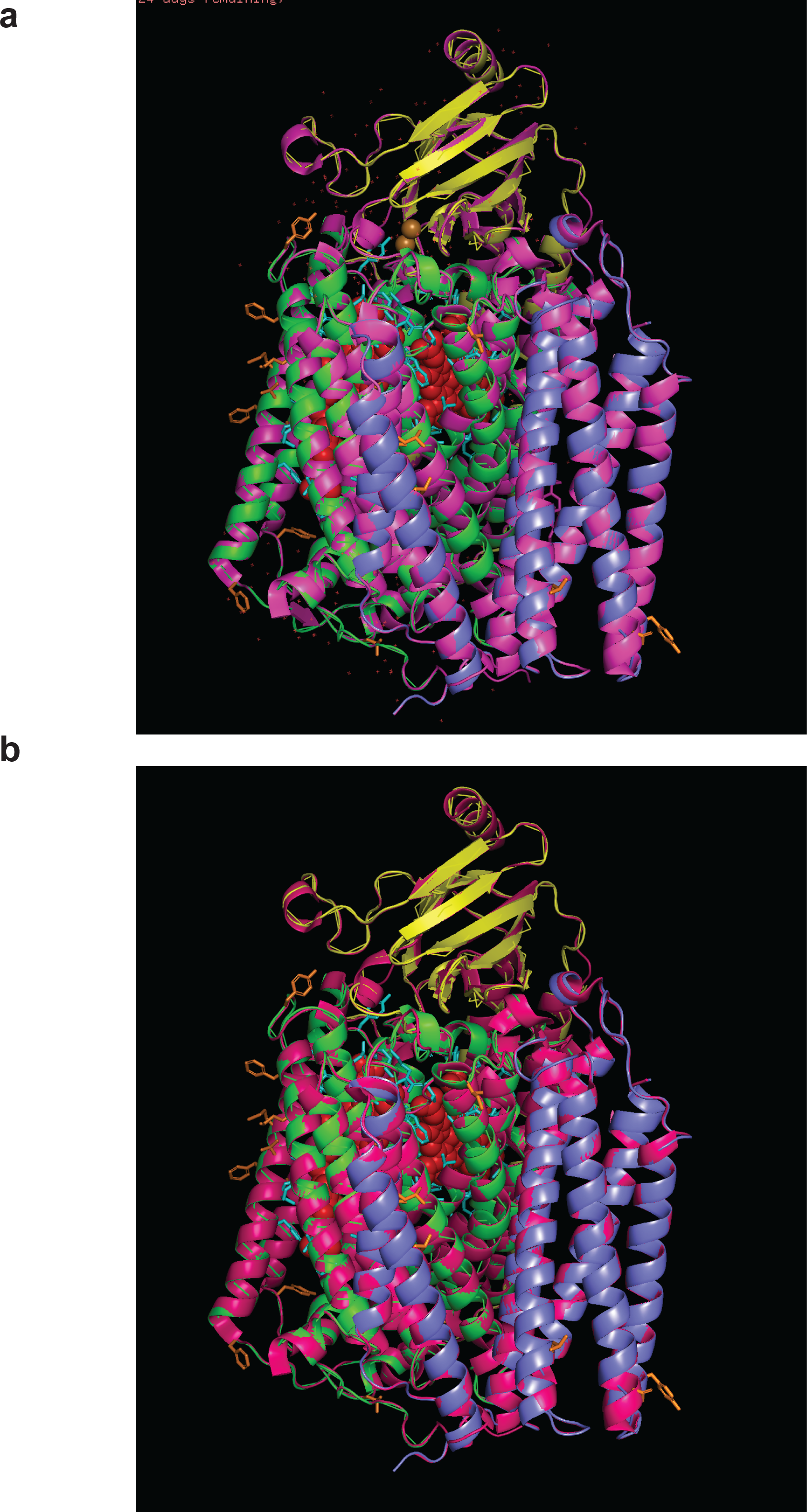
Superimposition of two bovine crystal structures with homology model of *H. mephisto* COX1, COX2, and COX3. **A.** Overlay of bovine structure 3abm with the homology model of *H. mephisto*. (Note: 3abm was used as the structural template for the homology model of *H. mephisto*.) **B.** Overlay of bovine structure 7coh with the homology structure of *H. mephisto.* For all panels, *H. mephisto* COX1 = green, COX2= yellow, and COX3=blue, and the bovine structure is pink.

**Figure S3.**
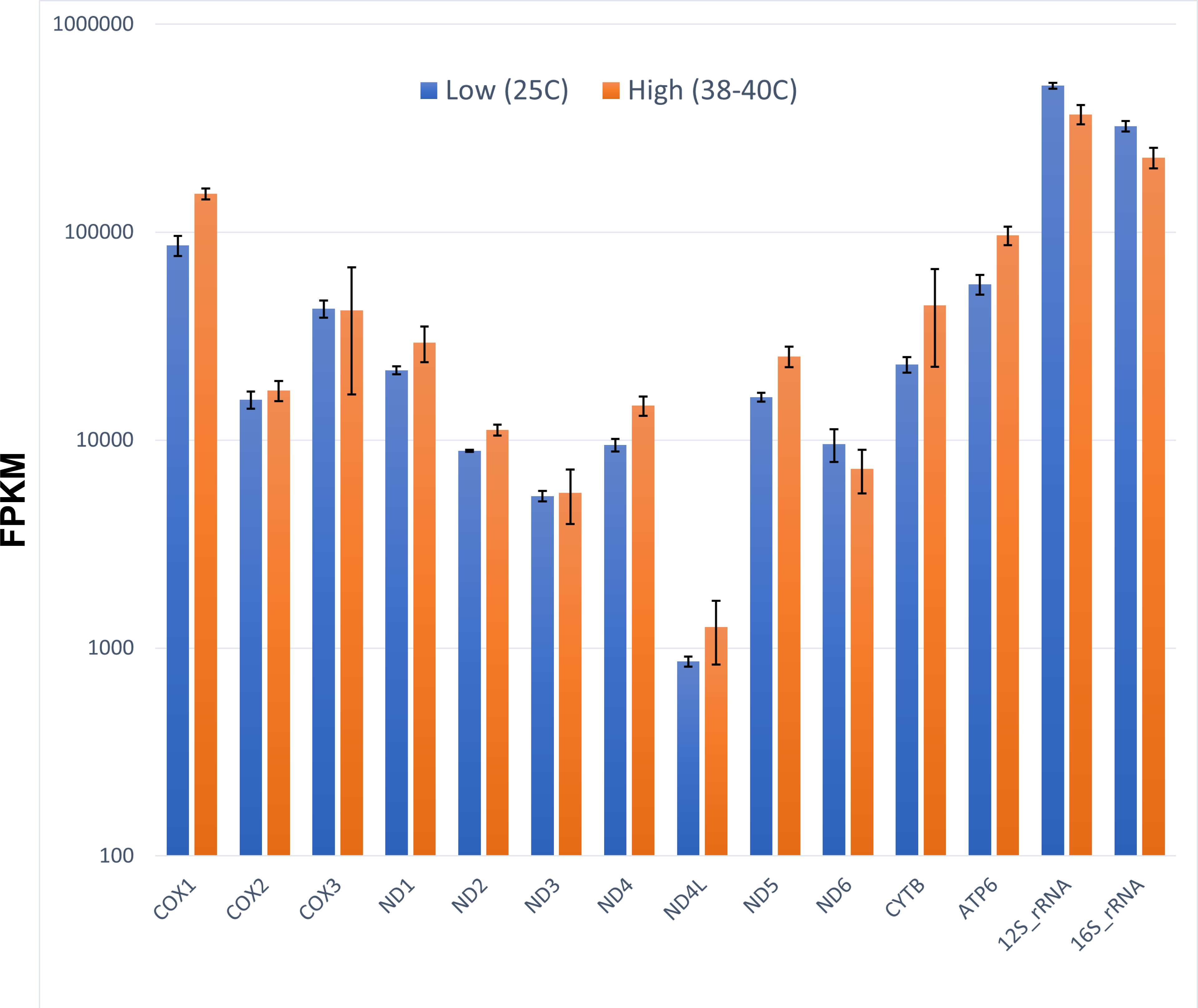
RNAseq expression of mitochondrial protein-coding genes at 25°C and 38-40°C. All temperature-driven changes are less than 2-fold. Shown is average FPKM, Fragments per Kb per Million mapped reads. Error bars indicate plus or minus one standard deviation.

**Figure S4.**
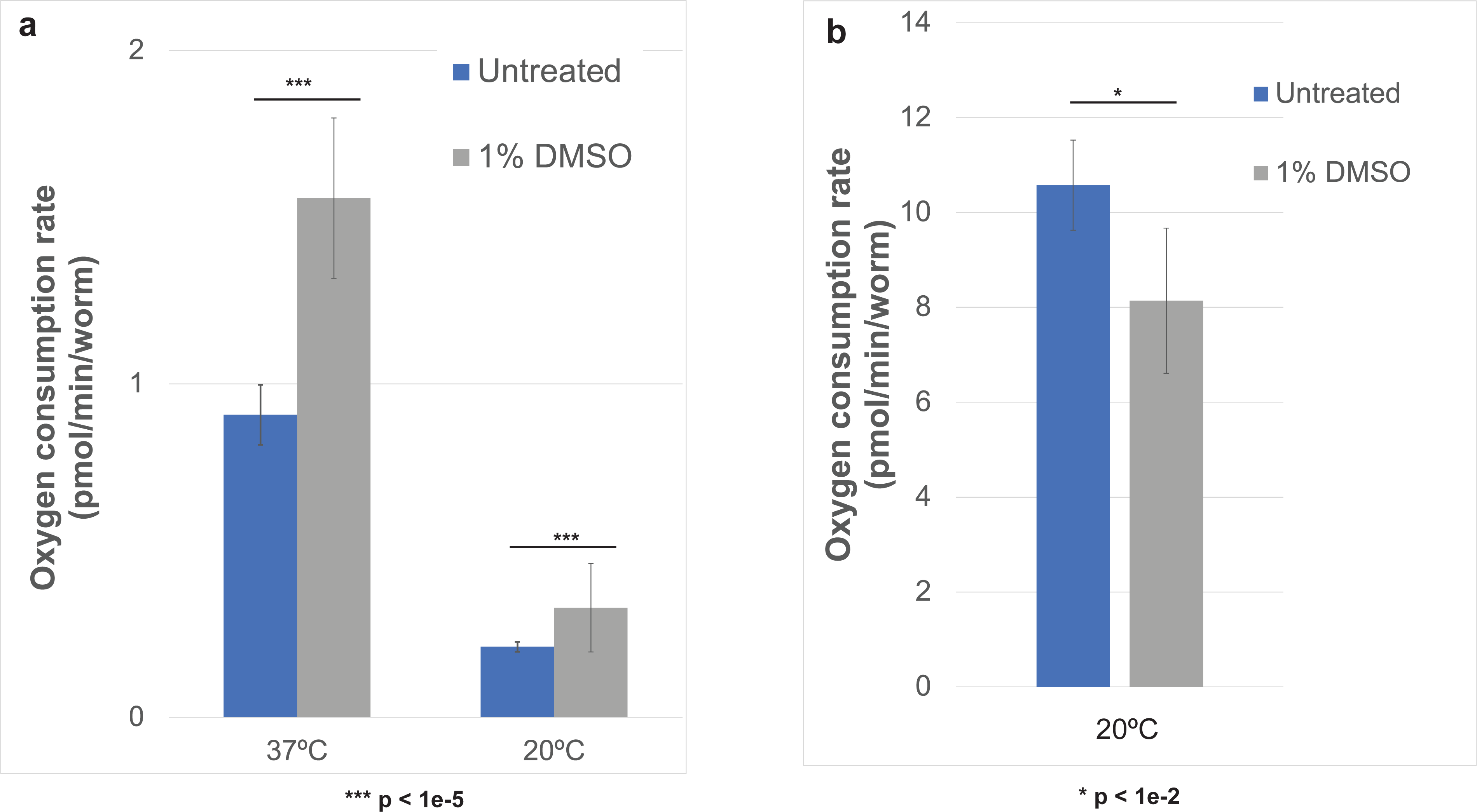
DMSO effect on oxygen consumption rate of a) *H. mephisto* and b) *C. elegans*. Shown is the mean with error bars representing standard error of the mean. p-values calculated in panel (a) by pairwise comparisons using Wilcoxon rank sum exact test and (b) by Student’s two-tailed T-test for samples of unequal variance.

## Materials and Methods

### Assembly of *H. mephisto* mitochondrial genome

As a component of a prior sequencing effort (Weinstein et al., 2019), PacBio RSII data were generated from raw genomic DNA. The data generated by three RSII lanes were assembled using HGAP3 and polished with Quiver (using the SMRT Analysis Software v2.3.0 pipeline), and the resultant assembly contained a contig encoding the complete mitochondrial genome (unitig_62|quiver).

### Curation of *H. mephisto* mitochondrial genome

The *H. mephisto* mitochondrial genome was aligned with that of *B. xylophilus* (AP017463.1), *P. redivivus* (AP017464.1), and *H. gingivalis* (KM192363.1) using MAFT v 7.017. Based on the clade IV mtDNA alignment and the results of MITOS WebServer v. 2.0^46^, annotations were manually curated to represent all coding sequences, two ribosomal RNA molecules, and 22 transfer RNAs, and an AT-rich control region.

### Assembly of *Diploscapter pachys* mitochondrial genome

*D. pachys* DNA was isolated without mtDNA enrichment and sequenced with Nanopore (version X). The genome was assembled from Nanopore reads by flye 2.8.1 and polished with Pilon 1.23. Pilon did not make any more corrections after iteration 3. Four indels were found by manual inspection of Illumina alignments, all of them in runs of Ts and within coding regions. These were then manually corrected to produce the current version of the D. pachys mitochondrial genome (13393 bps).

### Phylogenetic Analysis

For the catenated protein tree, all coding sequences concatenated in the following gene order: COX1, COX2, ND3, ND5, ND6, ND4L, ND1, ATP6, ND2, CYTB, COX3, and ND4. Concatenated amino acid sequences were aligned with MAFT v.7.4.90, and the ML tree constructed using IQ-Tree^47^ 1.6.12 webserver, allowing automatic detection of optimal substitution model (which for this dataset was mtZOA+F+I+G4). The Bayesian tree was constructed using Mr. Baye’s 3.2.6 from the same alignment with the equalin amino acid rate matrix and the invgamma rate parameter.

### dN/dS Analysis Using PAML’s Branch-Site Model

In this method individual branches are tested separately and a likelihood ratio test yields a p-value for the probability positive selection has occurred on the lineage; for positive branches specific amino acid codons are identified as selection sites^18^.To correct for multiple hypothesis testing, a simple Bonferroni correction was performed for each gene.

The nucleotide sequence for each protein-encoding gene was extracted and catenated together for each species. The resulting sequences were translation-aligned using Geneious Prime v.8.1.9 and all gaps were excised while maintaining coding frames of all sequences. Codeml of PAML v.4.9 was used to estimate branch site dN/dS, ⍵ for sequence alignments for the following genes: COX1, COX2, COX3, and NADH4. The branching clades for PAML were provided as the IQ-Tree ML phylogenetic tree as described above . In Codeml, branch site ⍵ was estimated using runmode = 0, seqtype = 1, model = 2, and NSsites = 2 in the control file. To enable likelihood ratio testing, every branch tested was run under two competing models: a neutral or relaxed selection model (fix_omega =1) and a positive selection model (fix_omega = 0). P-values were evaluated from the two likelihood values produced from the runs as follows. First we generated a LRT statistic, *2Δl.* This statistic was compared against *χ*^2^ tables with two degree of freedom (df=2). Thus, the critical value was set to 5.991 for 5% significance and 9.210 for 1% significance. To correct for multiple testing, we performed a Bonferroni correction: the raw p-values multiplied by the number of tests run in a single tree.

### Calculation of pairwise dS values

The Codeml package within PAML v. 4.9j was run in a pairwise mode as described^18^ and in the user documentation. Specifically, a pairwise codon alignment was provided in phylip format as a user tree, and the runmode variable was set to -2 (pairwise), seqtype = 1 (codons), model=1, and icode = 4 (for invertebrate mitochondrial genome translation table).

### COX Subunits Homology Modeling

SWISS-MODEL was accessed via the Expasy web server, which identified bovine crystal structure 3abm as a suitable template for modeling. *H. mephisto* COX1, COX2, and COX3 were provided for modeling. The resultant model displayed GMQE of 0.82 and QMEANDisCo of 0.75 +/- 0.05. The model was visualized in PyMol v. 2.5.1 and was virtually superimposable with two bovine structures 3abm (the template) and 7coh with RMSD of 0.142 (5880 of 5880 atoms) and 0.174 (5845 of 5845 atoms) angstrom, respectively. The H- and K-pathways were marked using the amino acid residues from^48^.

### Nematode culture, TMRE and sodium azide

The TMRE plates were prepared from standard, 60mm OP50-seeded NGM plates ^49^ supplemented with 500μL of 4μM TMRE dissolved in M9, evenly spread and allowed to soak into the plate and dry for 4-6 hours prior to adding nematodes. (Preparation of the 4μM stock was by 1:1000 dilution from a 4mM TMRE-DMSO stock. A control DMSO plate was prepared and used in parallel to control for autofluorescence. All images of DMSO-only nematodes were blank so are not shown).

Bleach-synchronized, starved L1 hatchling larvae of both *C. elegans* and *H. mephisto* were plated on NGM (no TMRE) and cultured at either 25°C (*C. elegans*) or 37°C (*H. mephisto*) for two days to reach adulthood. Worms were then washed to TMRE plates and cultured for 24 hours at the temperature for which the assay is to be conducted (20°C, 25°C, or 37°C), followed by destaining of TMRE for 1hr in 100μL M9, also at the assay temperature. Sodium azide (VWR Catalog # TS19038-0050) was used at 25mM final concentration and was added to appropriate samples during the 1hr TMRE destain period. All nematodes were paralyzed in a final concentration of 20mM Levamisole (AmBeed Cat # A121733-1G) immediately prior to imaging. For imaging, slides were made with 2% agarose pads in M9 and sealed with clear nail polish. Imaging was performed on an Olympus FV1200 scanning confocal microscope and 60x oil immersion objective (600x total magnification) focused on the body-wall muscle just behind the head area for each worm. Identical non-saturating laser settings were used on all images to ensure comparable, quantitative results, and images were processed using ImageJ v. 1.53t to determine the intensity of the spots per unit area, for a total of 40,240 analyzed regions. Statistical analysis was performed by two-way ANOVA and Tukey’s HSD post-hoc test in R version 4.2.1.

### Seahorse Analysis of oxygen consumption rates

A Seahorse HS Mini device was used for all Oxygen Consumption Rate (OCR) analysis, following nematode protocols as previously described^26,27^. We found that the Seahorse XFp fluxPak (Agilent 103022-100) is essential–not the HS Mini plates, which have a ring structure on the bottom of the microchamber that often excludes worms from the measurements. The XFp plates have a characteristic 3 dots visible in the microchamber but work much better.

To measure maximal respiration we used Carbonyl cyanide-p-trifluoromethoxy-phenylhydrazone (FCCP), a potent decoupling agent of the inner mitochondrial membrane. We tested a variety of concentrations and found maximal response at 40μM (final concentration) in both *C. elegans* and *H. mephisto.* Consistent with previous findings^26^, we observed that high (1%) DMSO final concentration was required for FCCP function; whether this is due to improved drug solubility or absorption by the nematodes is unclear but lower DMSO (<0.1%) yielded no response. Thus all analyses were performed in 1% DMSO final concentration and we measured a 1% DMSO control. By using FCCP and sodium azide (final concentration 25mM, soluble and functional in M9) we were able to measure basal, maximal, and spare capacity. Another compound, Dicyclohexylcarbodiimide (DCCD) has been shown to work in *C. elegans* to inhibit ATP synthase and thus to reveal ATP dependent oxygen consumption^26^; while we were able to replicate this, we found the drug had no effect on *H. mephisto* so we omitted it from our analysis.

We found that 11-13 *C. elegans* worms per well was optimal. Because *H. mephisto* are smaller we found that 60-80 worms per well was optimum. In all cases nematodes were visually counted under a light microscope before (*H. mephisto*) or before and after (*C. elegans*) a run. Because Seahorse analysis of nematodes is noisy^26^ we followed recommendations to allow many measurement cycles and take averages^26^. Therefore, one run consisted of 8 basal untreated measurements (prior to drug injection), 20 measurements after FCCP addition (or 1% DMSO control) and then 4 measurements after adding sodium azide addition. Each measurement cycle consisted of a 2 min mix, 2 min wait and 3 min measure period. We always performed 3 wells of FCCP injection and 3 wells of 1% DMSO injection, with two designated background wells.

Because DMSO had a strong effect on *H. mephisto* and *C. elegans* OCR (Figure S4), we used the 1% DMSO as basal measures in Figure 6e and 6f. (Untreated measurements for Figure S4 were taken from measurements 5-8 prior to any drug injections). We found that optimal measures for maximal responses were different between species, with *H. mephisto* peaking and dropping off more dramatically than *C. elegans*. Thus, we used four measurements, 10-13 for *H. mephisto*, whether of 1% DMSO controls (which we count as basal respiration in Figure 6e) or FCCP (for maximal respiration). For *C. elegans* we used measurements 13-16, in accord with Luz et al., 2015^26^, for the FCCP or 1% DMSO basal measure (Figure 6f).

Once we had obtained the data from the runs, we normalized OCR to pmol / min / worm and performed the following calculations following Agilent Seahorse guidelines: basal is 1% DMSO OCR minus the non-mitochondrial oxygen consumption (OCR after sodium azide). Maximal respiration is FCCP OCR minus non-mitochondrial OCR (sodium azide). Spare capacity is FCCP OCR minus 1% DMSO OCR.

### RNAseq data analysis for mitochondrial expression

Previously sequenced RNA-seq data ^2^ were mapped to the mitochondrial genome with HISAT2 2.2.1 and Stringtie 2.2.1 and Ballgown 2.26.0 installed and run in R 4.1.0. Differential expression was assessed with the stattest() function of Ballgown.

### Hypoxia culture and survival assessment

Nematodes were grown in anaerobic chambers. We seeded NGM plates with synchronized *C. elegans* or *H. mephisto* L1 larvae from overnight hatching in M9 (at 20°C for *C. elegans* and 37°C for *H. mephisto*), allowing a few minutes to dry, and then placing the plates into 2L chambers (Cat # 260002) with BD GasPak EZ anaerobe container system sachets (Cat # 260678) to create an anaerobic environment. Nematodes were cultured at either 20°C or 37°C depending on the species in standard incubators, for the indicated time (2 or 9 days) before opening and measuring viability. Survival was counted by washing the worms off the plates and exposing them to Sytox Orange dye (Thermo Fisher Cat # S11368), diluted 1:1000 in M9, for 15 minutes. Worms were then collected by centrifugation (400g for 2 minutes), pipetted onto 2% agarose-M9 pads, coverslipped, and sealed with clear nail polish prior to measurement of the dead animals using an Olympus BX61 fluorescence microscope set to the TRITC channel. Worm lengths (live animals only) were measured at the same time using CellSense v. 2.3 software.

