## Supplemental Figures for "Evolution of a biological thermocouple by adaptation of cytochrome c oxidase in a subterrestrial metazoan"

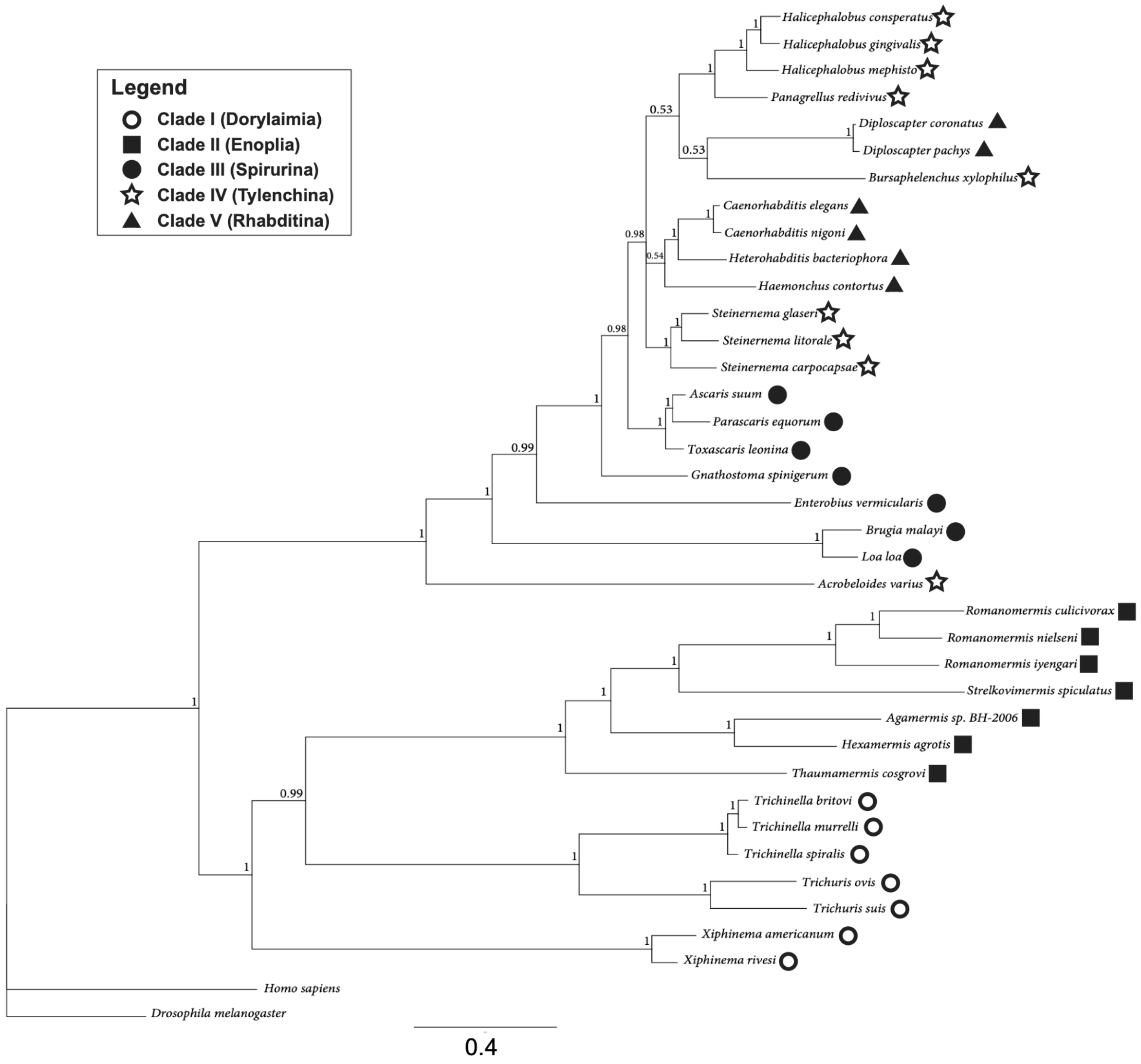


**Figure S1**. Bayesian catenated mitochondrial protein tree. Branches indicate posterior probabilities. Scale bar indicates substitutions per site.

a

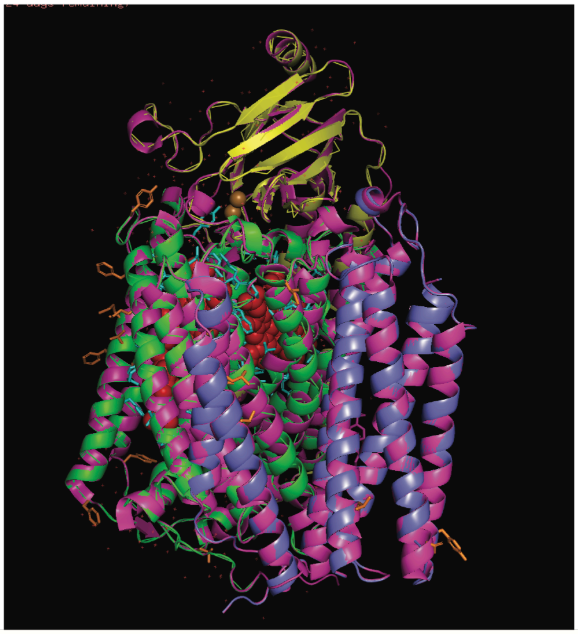

b

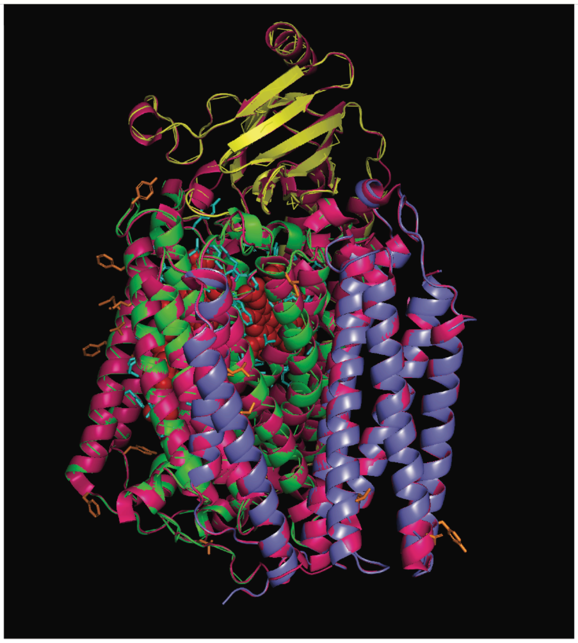

**Figure S2**. Superimposition of two bovine crystal structures with homology model of *H. mephisto* COX1, COX2, and COX3. **A.** Overlay of bovine structure 3abm with the homology model of *H. mephisto*. (Note: 3abm was used as the structural template for the homology model of *H. mephisto*.) **B.** Overlay of bovine structure 7coh with the homology structure of *H. mephisto.* For all panels, *H. mephisto* COX1 = green, COX2= yellow, and COX3=blue, and the bovine structure is pink.

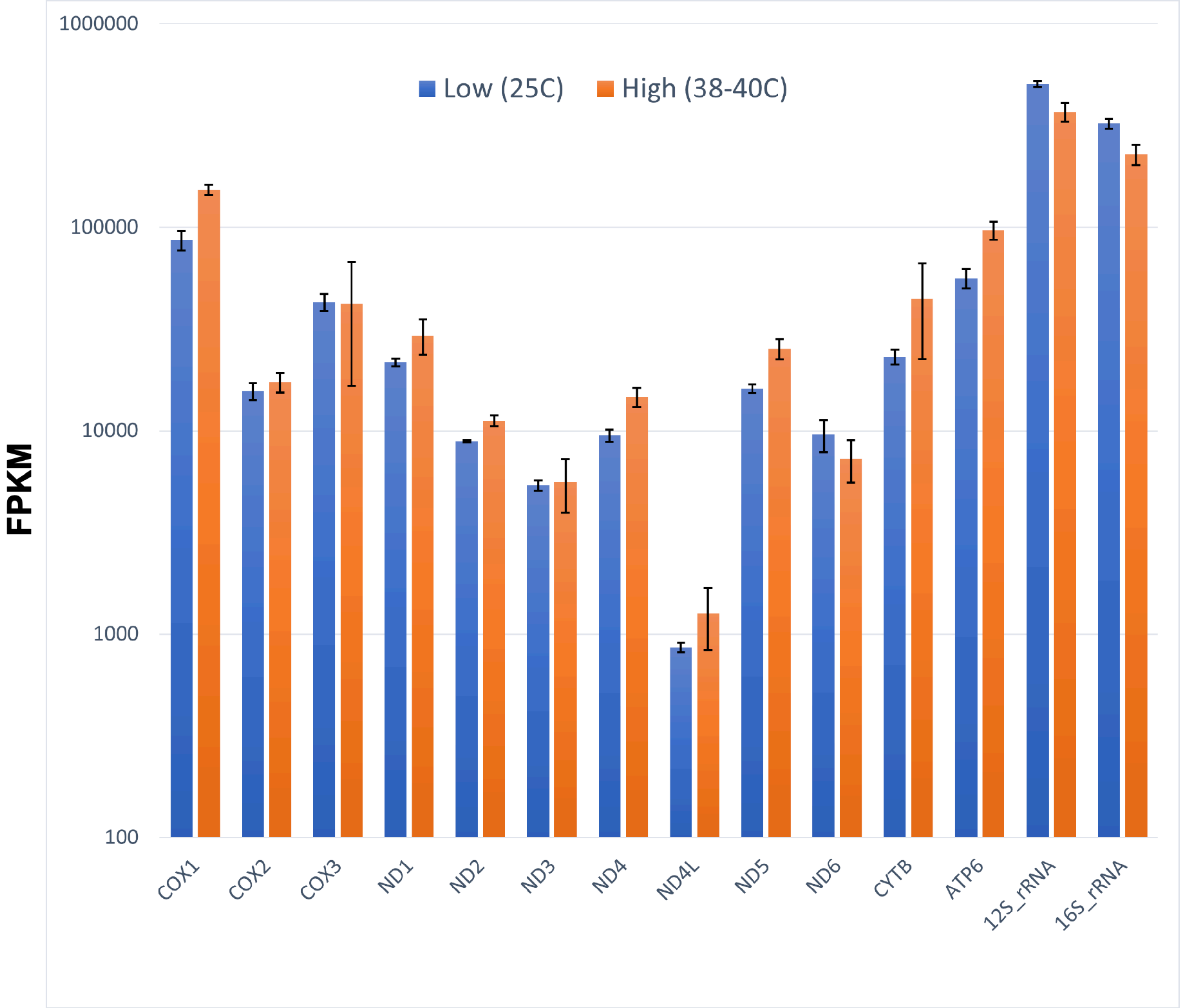


**Figure S3**. RNAseq expression of mitochondrial protein-coding genes at 25°C and 38-40°C. All temperature-driven changes are less than 2-fold. Shown is average FPKM, Fragments per Kb per Million mapped reads. Error bars indicate plus or minus one standard deviation.

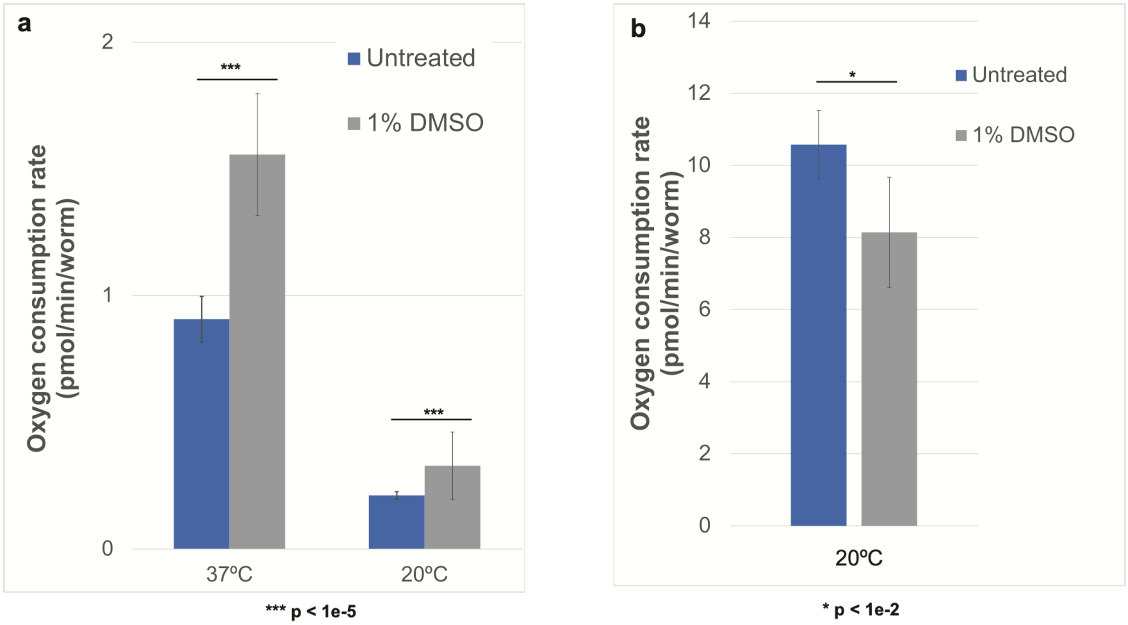


**Figure S4.** DMSO effect on oxygen consumption rate of a) *H. mephisto* and b) *C. elegans*. Shown is the mean with error bars representing standard error of the mean. p-values calculated in panel (a) by pairwise comparisons using Wilcoxon rank sum exact test and (b) by Student's two-tailed T-test for samples of unequal variance.
